# Efficient Federated Learning for distributed NeuroImaging Data

**DOI:** 10.1101/2024.05.14.594167

**Authors:** Bishal Thapaliya, Riyasat Ohib, Eloy Geenjar, Jingyu Liu, Vince Calhoun, Sergey Plis

**Affiliations:** TReNDS Center, Atlanta, Georgia, USA; Georgia Institute of Technology, Atlanta, Georgia, USA; Georgia State University, Atlanta, Georgia, USA

**Keywords:** Efficient Federated Learning, Neuro-imaging, Sparsity

## Abstract

Recent advancements in neuroimaging have led to greater data sharing among the scientific community. However, institutions frequently maintain control over their data, citing concerns related to research culture, privacy, and accountability. This creates a demand for innovative tools capable of analyzing amalgamated datasets without the need to transfer actual data between entities. To address this challenge, we propose a decentralized sparse federated learning (FL) strategy. This approach emphasizes local training of sparse models to facilitate efficient communication within such frameworks. By capitalizing on model sparsity and selectively sharing parameters between client sites during the training phase, our method significantly lowers communication overheads. This advantage becomes increasingly pronounced when dealing with larger models and accommodating the diverse resource capabilities of various sites. We demonstrate the effectiveness of our approach through the application to the Adolescent Brain Cognitive Development (ABCD) dataset.

## 1 INTRODUCTION

Deep learning has transformed fields like computer vision, natural language processing, and is also starting to transform the field of neuroimaging. As deep learning models grow, distributed and collaborative training becomes essential, especially when sensitive data is spread across distant sites. Collaborative MRI data analysis offers profound insights, allowing researchers to utilize data beyond a study’s original scope. As MRI scans are often preserved, vast amounts of data accumulate across decentralized research sites. Technological advancements have increased data complexity and reduced costs, prompting researchers to harness data from various groups for larger sample sizes and revealing significant features while preserving the privacy of the data.

Training models on more data, while preserving data privacy is thus crucial. Aggregating data from different sources to a central server for training can however expose this sensitive information, raising ethical concerns. Federated learning (FL) addresses this by allowing devices or organizations to train models locally and share training details without sharing the actual data.

In federated learning (FL), a central server coordinates training, and client sites communicate only model parameters, keeping local data private. In the decentralized setting, the server usually doesn’t exist and clients train a model collaboratively among themselves. However, challenges arise due to data’s statistical heterogeneity, limited communication bandwidth, and computational costs. In this work, we focus on addressing the communication efficiency inherent in distributed federated neuro-imaging pipelines through the training of sparse models at local sites.

### 1.1 Federated Learning

Federated Learning distinguishes itself from traditional distributed learning approaches, such as those discussed by McMahan et al. (2017), through its unique characteristics:

#### 1 Non-IID Data

The training data across different clients are not identically distributed, which means that the data at each local site may not accurately represent the overall population distribution.

#### 2 Unbalanced Data

The amount of data varies significantly across clients, leading to imbalances in data representation.

#### 3 Massive Distribution

Often, the number of clients exceeds the average number of samples per client, illustrating the scale of distribution.

#### 4 Limited Communication

Communication is infrequent, either among clients in a decentralized setting or between clients and the server in a centralized setting, due to slow and expensive connections.

One of the main focuses of this work is to reduce the communication costs between the server and clients in a centralized setting or among clients in a decentralized setting when dealing with non-IID and unbalanced data. This is achieved by identifying a sub-network based on the data distributions at each local site and transmitting only the parameters of this sub-network in each communication round *r*. In each round, a fixed set of 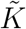 clients is sampled from all *K* clients, and federated training continues on the selected sub-network of those clients. The general federated optimization problem encountered is detailed next.

### 1.2 Federated Optimization Problem

In the general federated learning (FL) setting, a central server tries to find a global statistical model by periodically communicating with a set of clients. The federated averaging algorithm proposed by McMahan et al. (2017); Konečný et al. (2016); Bonawitz et al. (2019) is applicable to any finite sum objective of the form

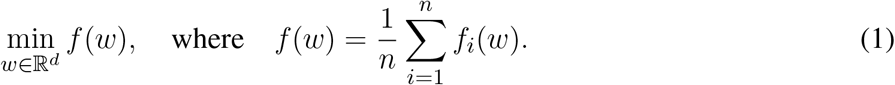

In a typical machine learning problem, the objective function *f*_*i*_(*w*) = *𝓁*(*x*_*i*_, *y*_*i*_; *w*) is encountered, where the *i*^th^ term in the sum is the loss of the network prediction on a sample (*x*_*i*_, *y*_*i*_) made by a model with parameter *w*. We assume that the data is partitioned over a total of *K* clients, with *𝒫*_*k*_ denoting the set of indices of the samples on client *k*, and *n*_*k*_ = |*𝒫*_*k*_|. Thus, the objective in (1) can be re-written as

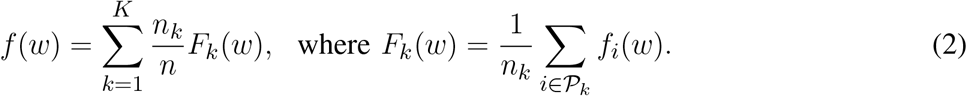

In the typical distributed optimization setting, the IID assumption is made, which says the following: if the partition *𝒫*_*k*_ was created by distributing the training data over the set of clients uniformly at random, then we would have 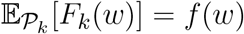, where the expectation is over the set of examples assigned to a fixed client *k*. In this work, we consider the non-IID setting where this does not hold and *F*_*k*_ could be an arbitrarily bad approximation to *f*.

When designing an FL training paradigm, a set of core considerations have to be made to maintain data privacy and address *statistical* or *objective* heterogeneity due to the differences in client data and resource constraints at the client sites. A range of work tries to address the issue of heterogeneous non-IID data McMahan et al. (2016); Kulkarni et al. (2020), however, some research also suggests that deterioration in accuracy in the FL non-IID setting is almost inevitable Zhao et al. (2018).

### 1.3 Federated Learning in Neuro-imaging

Over the past decade, the field of neuroimaging has strongly embraced data sharing, open-source software, and collaboration across multiple sites. This shift is largely attributed to the high costs and time demands associated with neuroimaging data collection Landis et al. (2016); Rootes-Murdy et al. (2022). By pooling data from different sources, researchers can explore findings that extend beyond the initial scope of individual studies Poldrack et al. (2013). The practice of sharing data enhances the robustness of research through larger sample sizes and the replication of results, offering significant benefits for neuroimaging studies. Increasing the sample size not only makes predictions more reliable but also ensures the reliability and validity of research findings, thereby preventing data manipulation and fabrication Ming et al. (2017); Tenopir et al. (2011). Furthermore, aggregating data can lead to a more diverse sample by combining otherwise similar datasets, thus reflecting a broader range of social health determinants for more comprehensive results Laird (2021). Additionally, reusing data can significantly reduce research costs Milham et al. (2018).

Federated Learning (FL) is increasingly recognized as a transformative approach in healthcare and neuro-imaging. In the realm of biomedical imaging, FL has been applied to a variety of tasks. These include whole-brain segmentation from MRI T1 scans Roy et al. (2019), segmentation of brain tumors Sheller et al. (2019); Li et al. (2019), multi-site fMRI classification, and the identification of disease biomarkers Li et al. (2020b). COINSTAC Plis et al. (2016a) offers a privacy-focused distributed data processing framework specifically designed for brain imaging showcasing FL’s role in enhancing privacy and efficiency in healthcare data analysis. Additionally, it has been utilized in discovering brain structural relationships across various diseases and clinical cohorts through federated dimensionality reduction from shape features Silva et al. (2019).

### 1.4 Efficiency in Federated Learning

The primary objective of *model pruning* is to identify subnetworks within larger architectures by selectively removing connections. This technique holds considerable appeal for various reasons, particularly for real-time applications on resource-constrained edge devices, which are prevalent in federated learning (FL) and collaborative learning scenarios. Pruning large networks can significantly alleviate the computational demands of inference Elsen et al. (2020) or hardware tailored to exploit sparsity Cerebras (2019); Pool et al. (2021). More recently, the *lottery ticket hypothesis* has emerged Frankle and Carbin (2019), suggesting the existence of subnetworks within densely connected networks. These subnetworks, when trained independently from scratch, can attain comparable accuracy to fully trained dense networks Frankle and Carbin (2019), revitalizing the field of sparse deep learning Renda et al. (2020); Chen et al. (2020). This resurgence of interest has also extended into sparse reinforcement learning (RL) Arnob et al. (2021); Sokar et al. (2021). Pruning techniques in deep learning can broadly be categorized into three groups: methods that induce sparsity before training and during initialization Lee et al. (2018); Wang et al. (2020); Tanaka et al. (2020), during training Zhu and Gupta (2018); Ma et al. (2019); Yang et al. (2019); Ohib et al. (2022), and post-training Han et al. (2015); Frankle et al. (2021).

For pruning in the FL setting, using a *Lottery Ticket* like approach would result in immense inefficiency in communication. Such methods Frankle and Carbin (2019); Bibikar et al. (2022) usually require costly pruning and retraining cycles, often training and pruning multiple times to achieve the desired accuracy vs sparsity trade-off. Relatively few research have leveraged pruning in the FL paradigm Li et al. (2020a, 2021); Jiang et al. (2022). In particular, with LotteryFL Li et al. (2020a) and PruneFL Jiang et al. (2022), clients need to send the full model to the server regularly resulting in higher bandwidth usage. Moreover, in Li et al. (2020a), each client trains a personalized mask to maximize the performance only on the local data. A few recent works Bibikar et al. (2022); Huang et al. (2022); Qiu et al. (2022); Li et al. (2020a) also attempted to leverage sparse training within the FL setting as well. In particular Li et al. (2020a) implemented randomly initialized sparse mask, FedDST Bibikar et al. (2022) built on the idea of RigL Evci et al. (2020) and mostly focussed on magnitude pruning on the server-side resulting in similar constraints and Ohib et al. (2023) uses sparse gradients to efficiently train in a federated learning setting. In this work, we try to alleviate these limitations which we discuss in the following section.

## 2 METHOD DESCRIPTION

In this section we present our proposed method. We first describe the process of discovering a sub-network *f* (***θ*** *⊙* **m**) within the full network *f* (***θ***), where ***θ*, m** ∈ ℝ^*d*^ with ∥**m**∥_0_ *< d*. To discover a performant sub-network an importance scoring metric is required, which we describe in section 2.1.1. Finally, we delineate our proposed method in section 2.2

### 2.1 Sub-network discovery

Given a dataset 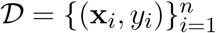 at a site *k*, the training of a neural network *f* parameterized by *θ* ∈ ℝ^*d*^ can be written as minimizing the following empirical risk:

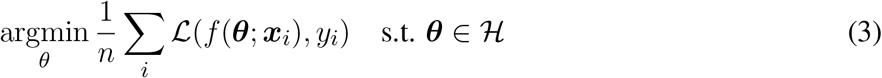

where ***θ*** ∈ ℝ^*d*^ and *ℒ* and *ℋ* are the loss function and the constraint set respectively.

In general, in unconstrained (standard) training the set of possible hypotheses is considered to be *ℋ* = ℝ^*d*^, where *d* is the model dimension. The objective is to minimize the empirical risk *ℒ* given a training set 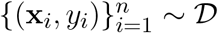 at the local client site *k*. Given access to the gradients of the empirical risk on a batch-wise basis, an optimization algorithm such as Stochastic Gradient Descent (SGD) is typically employed to achieve the specified objective. This process generates a series of parameter estimates, 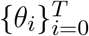, where *θ*_0_ represents the initial parameters and *θ*_*T*_ the final optimal parameters. A sub-network within this network is defined as a sparse version of this network with a mask **m** ∈ {0, 1}^|***θ***|^ that results in a masked network *f* (***θ*** *⊙* **m**; ***x***_*i*_). When aiming for a target sparsity level where *k < d*, the parameter pruning challenge entails ensuring that the final optimal parameters, *θ*_*T*_, have at most *k* non-zero elements, as denoted by the constraint ∥*θ*_*T*_ ∥_0_ *≤ k*. In many works, this sparsity constraint applies only to the final parameters and not to any intermediate parameter estimates. However, in this work we maintain this sparsity constraint throughout the entire training phase, that is throughout the entire evolution of *θ* from *θ*_0_ to *θ*_*T*_.

The goal of discovering sub-networks at initialization introduces additional constraints to the previously described framework by requiring that all parameter iterations fall within a predetermined subspace of *ℋ*. Specifically, the constraints seek to identify an initial set of parameters, *θ*_0_, that has no more than *k*_1_ non-zero elements (∥*θ*_0_∥_0_ *≤ k*_1_), and ensure that all intermediate parameter sets, *θ*_*i*_, belong to a subspace 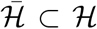 for all *i* in {1, …, *T* }, where 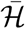 is the subspace of ℝ^*d*^ spanned by the natural basis vectors 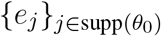. Here, supp(*θ*_0_) represents the support of *θ*_0_, or the set of indices corresponding to its non-zero entries. This approach not only specifies a sub-network at initialization with *k* parameters but also maintains its structure consistently throughout the training.

#### 2.1.1 Connection Importance Criterion

Lee et al. (2018) introduced a technique inspired by the saliency criterion originally proposed by Mozer and Smolensky (1988). They contributed an important insight, demonstrating that this criterion is remarkably effective in predicting the significance of each connection in a neural network at the initialization phase. The core concept revolves around retaining those parameters that, when altered, would have the most substantial effect on the loss function. This is operationalized by considering a binary vector *c* ∈ {0, 1}^*m*^ and utilizing the Hadamard product *⊙*. Consequently, SNIP calculates the sensitivity of connections based on this approach. For each connection *θ*_*j*_ in the network, the importance score for that connection is calculated as following:

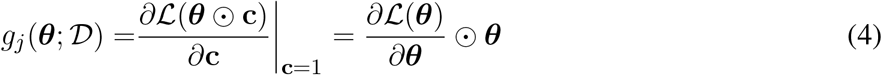

After determining *g*(*θ*), the parameters associated with the highest *k* magnitudes of |*g*_*j*_(*θ*; *D*)| are retained. Essentially, SNIP prioritizes weights that, regardless of their direction, are distant from the origin and yield large gradient values. It’s noteworthy that the objective of SNIP can be reformulated as follows De Jorge et al. (2020); Frankle et al. (2021):

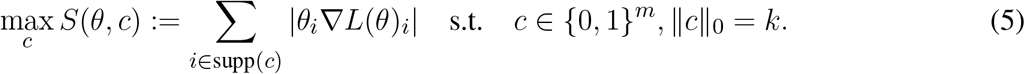

It is trivial to note that the optimal solution to the above problem can be obtained by selecting the indices corresponding to the top-*k* values of |*θ*_*i*_ *∇ℒ* (*θ*_*j*_)|.

### 2.2 Proposed Method

We propose a novel method for *efficient distributed sub-network discovery* for distributed neuroimaging and propose a method for training such sparse models or subnetworks in a communication efficient manner called *Sparse Federated Learning for Neuro-Imaging* or NeuroSFL with the goal of tackling communication inefficiency during decentralized federated learning with non-IID data distribution in the context of distributed neuro-imaging data. The proposed method initiates with the common initialization ***θ***_0_ at all the local client models. Next, importance scores *s*_*j*_ are calculated for each model parameter in the network based on the information from the imaging data available across all the client sites. At this stage, each client has a unique set of importance scores for their parameters in the local network *f* based on the local data available at that site similar to Lee et al. (2018); De Jorge et al. (2020). All the clients transmit these scores to each other and a mask **m** is created corresponding to the top-*k* % of the aggregated saliency scores:

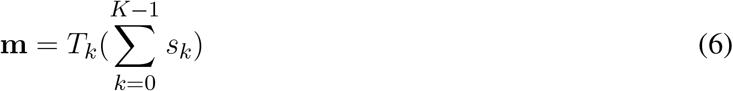

Where the *T*_*k*_ is the top-k operator. This mask is then used to train the model *f*_*k*_(***θ*** *⊙* **m**; ***x***) at site *k* on their local data (***x***, *y*) *∼ 𝒟*_*k*_.

For the federated training among a total of *K* clients, the clients are trained locally, and at the end of local training they share their trained parameters which are then averaged; we call this a *communication round*. At the start of this local training, each site *k* starts with the same initial model weights *θ*_0_ which at each site *k* is denoted as *θ*_*k*,0_ at training step *t* = 0 which are then masked with the generated saliency mask **m** to produce the common masked initialization 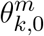 as follows:

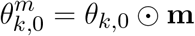

Next these models at each site *k* are trained on their local dataset (***x***, *y*) *∼ 𝒟*_*k*_.

The masked models *f* (*θ*_*k*,0_ *⊙* **m**) across all the sites are trained for a total of *T* communication rounds to arrive at the final weights *θ* _*k,T*_ at each local site. In each communication round *t*, only a random subset ℱ′ = {*f*_1_, *f*_2_, …, *f*_*K*′_ } of *K*′ clients where ℱ′ ⊆ ℱ the set of all clients, and *K*′ *≤ K* are trained on their local data (a way to avoid the *straggler effect* in the real world, wherein a large client group the update might be bottle-necked by the slowest, most resource-constrained client site). At the end of local training, the selected subset ℱ′ of the clients contribute to training the updated model 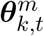, which is the starting weights for the next communication round. When sharing the updated weights only the weights corresponding to the 1’s in the binary mask **m** are shared among the clients and with the server, as only these weights are being trained and the rest of the weights are *zero-ed* out. This results in the gains in communication efficiency. To efficiently share the model weights, the clients only share their sparse masked weights 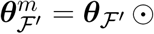 among the selected clients in ℱ′ using the compressed sparse row (CSR) encoding. The algorithm for the training process is delineated in Algorithm 1.

### 2.3 Iterative Importance Score Calculation: IterativeSNIP

In this section, we test the effectiveness of iterative-SNIP De Jorge et al. (2020), which is an iterative version of the application of saliency criterion in eq. (4). We briefly describe the iterative-SNIP next. We assume *k* to be the number of parameters to be preserved post pruning. Given that we have some pruning schedule (similar to learning rate schedule: linear, exponential etc.) to divide *k* into a set of natural numbers 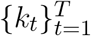 such that *k*_*t*_ *> k*_*t*+1_ and *k*_*T*_ = *k*. Now, given the binary masking variable ***c***_*t*_ corresponding to *k*_*t*_, the formulation of pruning from *k*_*t*_ to *k*_*t*+1_ can be made using the connection sensitivity (4) similar to De Jorge et al. (2020) as:

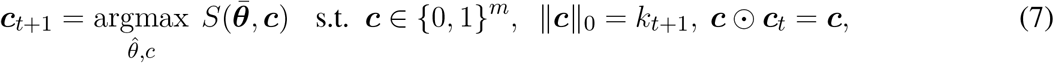

where 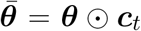. The constraint ***c*** *⊙* ***c***_*t*_ = ***c*** is added to ensure that no previously pruning parameter is re-activated. Assuming that the pruning schedule ensures a smooth transition from one topology to another (∥***c***_*t*_∥_0_ *≈* ∥***c*** _*t*+1_∥_0_) such that the *gradient approximation* 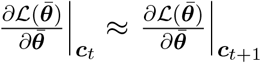 is valid, (7) can be approximated as solving (5) at 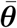

In the scenario where the schedule parameter is set to *T* = 1, the original SNIP saliency method is recovered. This is basically employing a *gradient approximation* approach between the initial dense network ***c***_0_ = **1** and the resulting mask ***c***. We conduct experiments with IterativeSNIP in the federated neuroimaging setting and present our findings in section 4.2.

#### Algorithm 1

NeuroSFL

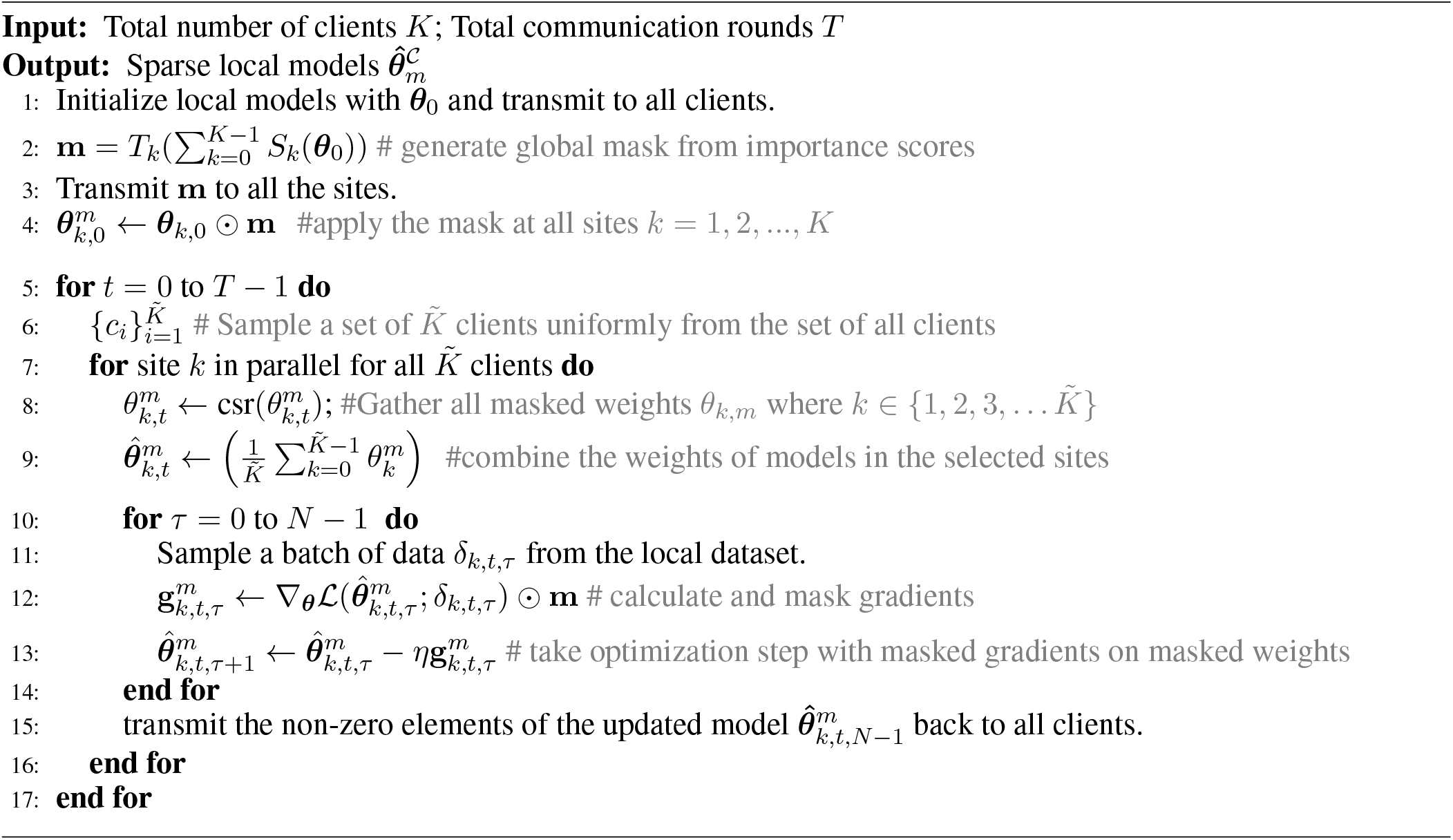

## 3 EXPERIMENTS

### 3.1 Dataset and non-IID partition

We evaluated NeuroSFL on the ABCD dataset. ABCD study is the largest long-term study of brain development and child health in the US. It recruited over 10 thousand children of 9 and 10 years old from 21 sites and followed them for 10 years with annual behavioral and cognitive assessments and biannual MRI scans Garavan et al. (2018). Along with multi-session brain MRI scans for structure and function, the ABCD study also includes key demographic information including gender, racial information, socio-economic backgrounds, cognitive development, and mental and physical health assessments of the subjects. The ABCD open-source dataset can be found on the National Institute of Mental Health Data Archive (NDA) https://nda.nih.gov/. In this study, we used data from the ABCD baseline, which contain 11,875 participants aged 9-10 years.

T1-weighted MRI images were preprocessed using the Statistical Parametric Mapping 12 (SPM12) software toolbox for registration, normalization, and tissue segmentation. Then the gray matter density maps were smoothed by a 6 mm^3^ Gaussian kernel, creating images with the dimensionality of (121,145,121) of voxels at Montreal Neuroimaging Institute (MNI) space with each voxel having dimensions of 1.5×1.5×1.5^3^ mm.

We simulated the heterogeneous data distributions across federated clients through the adoption of two distinct data partitioning strategies. We outline these strategies for generating non-IID data partitions with a comprehensive discussion in section 3.1.1.

#### 3.1.1 Generating non-IID data partition with Dirichlet Distribution

In this section, we provide the necessary background on generating non-identical data distribution in the client sites using the Dirichlet Distribution, specifically for the context of federated learning.

##### 3.1.1.1 non-IID data in FL

Federated Learning (FL), as introduced by McMahan and Ramage (2017), is a framework designed for training models on decentralized data while preserving privacy. It utilizes the Federated Averaging (FedAvg) algorithm where each device, or client, receives a model from a central server, performs stochastic gradient descent (SGD) on its local data, and sends the models back for aggregation. Unlike data-center training where data batches are often IID (independent and identically distributed), FL typically deals with non-IID data distributions across different clients. Hence, to evaluate federated learning it is crucial to not make the IID assumption and instead generate non-IID data among clients for evaluation Hsu et al. (2019).

##### 3.1.1.2 Generating non-IID data from Dirichlet Distribution

In this study, we assume that each client independently chooses training samples. These samples are classified into *N* distinct classes, with the distribution of class labels governed by a probability vector ***q***, which is non-negative and whose components sum to 1, that is, *q*_*i*_ *>* 0, *i* ∈ [1, *N* ] and ∥***q***∥_1_ = 1. For generating a group of non-identical clients, *q ∼* Dir(*α****p***) is drawn from the Dirichlet Distribution, with ***p*** characterizing a prior distribution over the *N* classes and *α* controls the degree of identicality among the existing clients and is known as the *concentration parameter*.

In this section, we generate a range of client data partitions from the Dirichlet distribution with a range of values for the concentration parameter *α* for exposition. In fig. 1, we generate a group of 10 balanced clients, each holding an equal number of total samples. Similar to Hsu et al. (2019) the prior distribution ***p*** is assumed to be uniform across all classes. For each client, given a concentration parameter *α*, we sample a ***q*** from Dir(*α*) and allocate the corresponding fraction of samples from each client to that client. fig. 1 illustrates the effect of the concentration parameter *α* on the class distribution drawn from the Dirichlet distribution on different clients, for the CIFAR-10 dataset. When *α → ∞*, identical class distribution is assigned to each classes. With decreasing *α*, more non-identicalness is introduced in the class distribution among the client population. At the other extreme with *α →* 0, each class only consists of one particular class.

**Figure 1.**
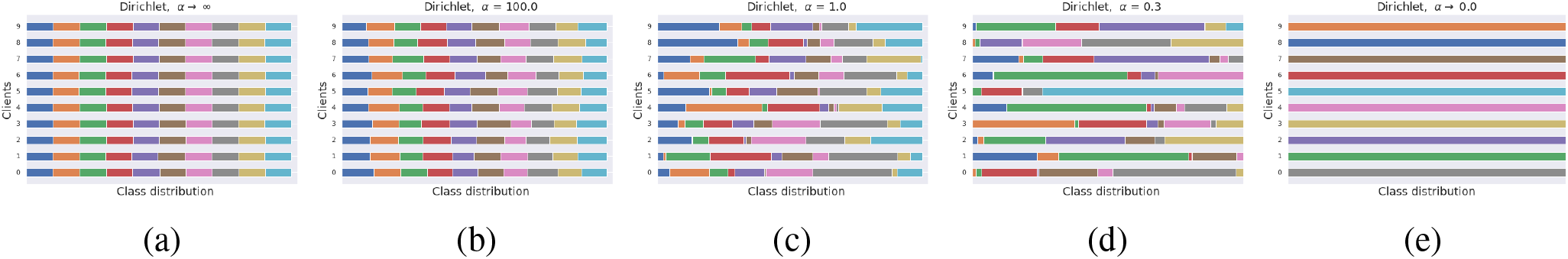
Generating non-identical client data partitions using the Dirichlet Distribution for the Cifar10 dataset among 10 clients. Distribution among classes is represented using different colors. (a) Dirichlet, *α → ∞* results in identical clients (b-d) Client distributions generated from Dirichlet distributions with different concentration parameters *α* (e) Dirichlet, *α →* 0.0 results in each client being assigned only 1 particular class.

### 3.2 Architecture, Hyperparameters and experimental details

Here we provide a comprehensive overview of the architecture, hyperparameters, and the experimental setup we use to evaluate our proposed NeuroSFL method on the neuroimaging Adolescent Brain Cognitive Development (ABCD) data. Our study focuses on the task of classifying a participant’s sex based on MRI scans, by employing a 3D variant of the well-known AlexNet model Krizhevsky et al. (2012). The 3D variant was referenced from Abrol et al. (2021), which has a specific channel configuration for the convolutional layers set as: 64C-128C-192C-192C-128C, where ‘C’ denotes channels.

We optimized the learning rate for this task through an exhaustive search ranging from LR=1 *×* 10^*−*3^ to 1 *×* 10^*−*6^, achieving a delicate balance between rapid convergence and fine-tuning during training. We employed a batch size of 32 and a learning rate decay factor of 0.998 was applied. We applied varying sparsity levels, ranging from 0%, 50%, 80%, 90%, and 95% to assess the overall performance. A split of 80/20 was used for training and testing for each individual site. Our training consists of 5 epochs with 200 communication rounds.

### 3.3 Baselines

We compared our method with both centralized and decentralized baselines. Centralized baseline includes FedAvg McMahan et al. (2017), FedAvg-FT Cheng et al. (2021) which are the standard dense baselines, and for the decentralized FL setting, we take the sparse Dis-PFL Dai et al. (2022).

In FedAvg McMahan et al. (2017), each client trains its local model using its local data, and then these local models are aggregated or averaged to update the global model. On the other hand, FedAvg-FT Cheng et al. (2021) extends the FedAvg algorithm by incorporating fine-tuning or transfer learning. Specifically, after the global model is trained using FedAvg, the global model is then fine-tuned or adapted using additional data from a central server or other external sources. This fine-tuning step allows the global model to adapt to new tasks or data distributions beyond what was initially learned from the federated learning process. We also compare with DisPFL Dai et al. (2022) with varying sparsity levels. DisPFL is a new sparse FL technique that randomly prunes each layer similar to Evci et al. (2020) and uses the prune and regrow method from that work as well, resulting in a dynamically sparse method.

In exploring the impact of using unique local masks instead of a global mask on the performance of FL, we established IndividualSNIP as a baseline, representing an approach where unique local masks are devised from the saliency criterion, and local models are trained based on these masks. Moreover, to isolate the impact of just using *global masking*, that is using the same random mask in all clients, instead of using different unique random masks at different sites we compare our method and competing methods against random global masking as well in fig. 2.

**Figure 2.**
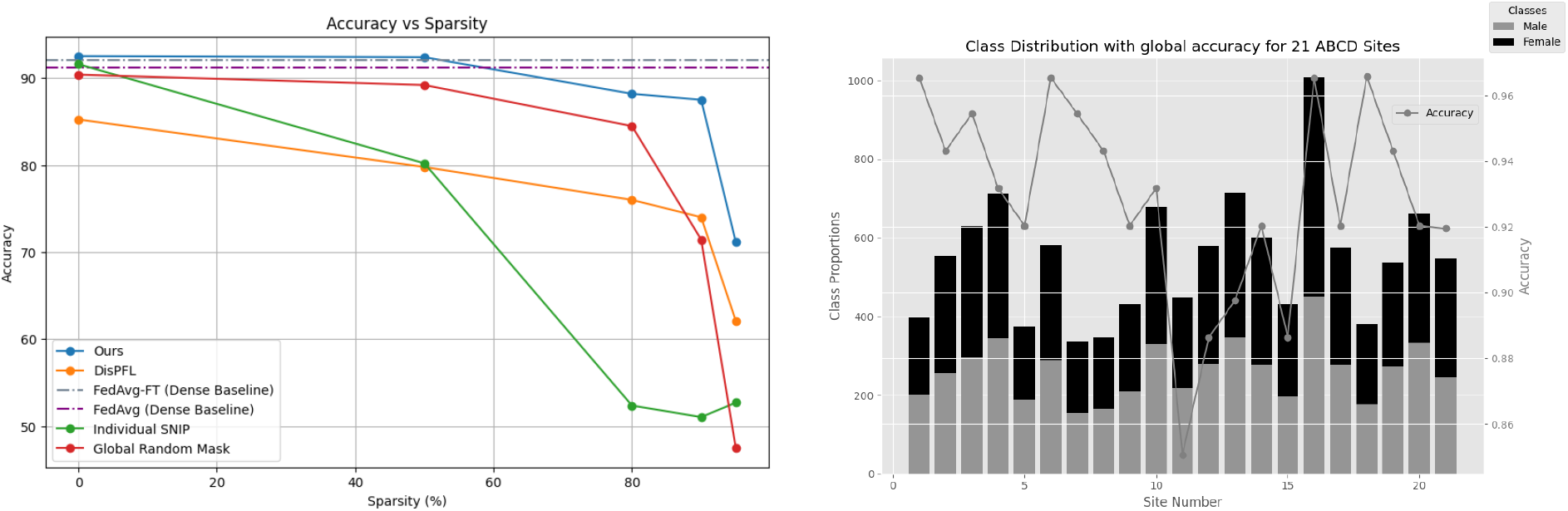
(a) Comparison of methods for gender classification using MRI Scans of ABCD dataset. (b) Gender differences in each of the 21 ABCD sites along with the performance of the global model on each site with 50% sparsity.

Additionally, with further exploration to investigate how different methods of model pruning and selection impact the performance of our approach, we further experiment with other techniques named IterSNIP and WeightedSNIP. IterSNIP builds upon the traditional SNIP method Lee et al. (2018) by incorporating multiple minibatches during the training process of mask generation. This approach aggregates saliency scores from these mini-batches to generate a comprehensive and robust pruning mask. Conversely, WeightedSNIP adopts a different strategy, deriving a global mask through a weighted average of saliency scores based on the frequency of data at each site, and assigning importance levels to individual sites based on the amount of data at sites.

## 4 RESULTS AND DISCUSSION

### 4.1 Effect of varying sparsity levels

We first explore the effect of sparsity on IID data in section 4.1.1 and then explore the efficacy of NeuroSFL on non-IID data in asection 4.1.2.

#### 4.1.1 Effect of varying sparsity levels on IID data

The performance of various methods across different sparsity levels was evaluated, as presented in Table 1, and visually presented in Figure 2. Sparse baselines, including Ours (*NeuroSFL*), IndividualSNIP, DisPFL Dai et al. (2022), and Global Random Mask, were compared against dense baselines such as FedAvg-FT Cheng et al. (2021) and FedAvg McMahan et al. (2017). Notably, our proposed *NeuroSFL*, exhibited robust performance across varying sparsity levels, achieving an accuracy of 92.52% at 0% sparsity and maintaining high accuracy even at higher sparsity levels, with 71.18% accuracy at 95% sparsity. In comparison, IndividualSNIP demonstrated decreasing accuracy as sparsity increased, with a significant drop to 52.70% at 95% sparsity. This is in line with expectation as individual-SNIP only incorporates the saliency scores from a single site at random and does not incorporate information from the datasets at all the participating cites.

**Table 1.**
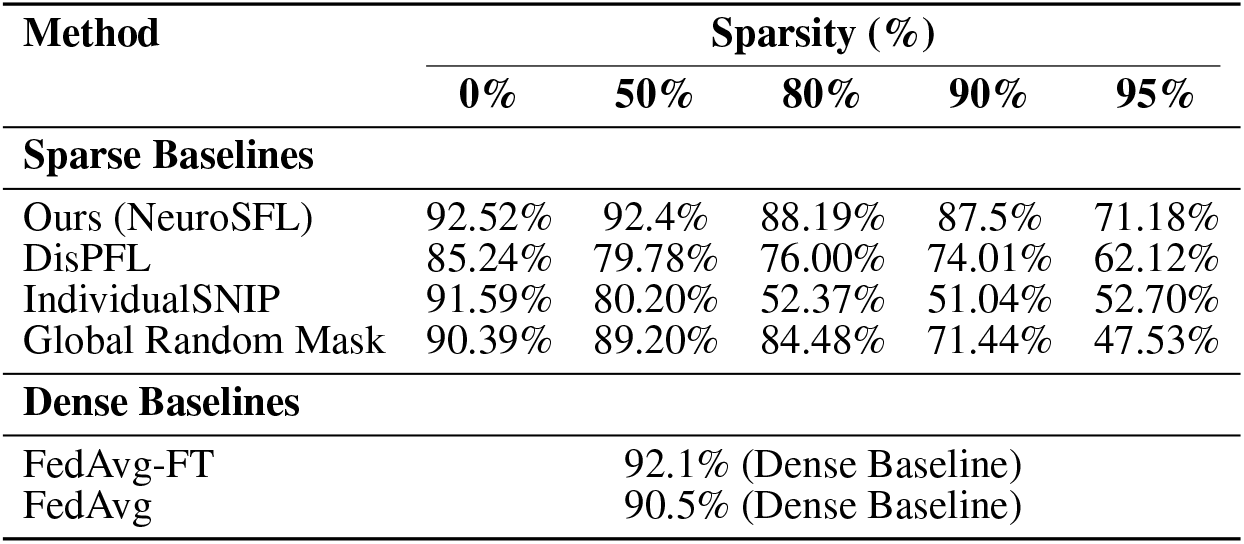
Performance comparison of different methods and sparsity levels. Sparse baselines include Ours-(NeuroSFL), DistPFL, IndividualSNIP, and Global Random Mask. Dense baselines include FedAvg and FedAvg-FT.

Moreover, in contrast to NeuroSFL, DisPFL, and Global Random Mask also showcased diminishing accuracy with increasing sparsity, highlighting the effectiveness of our proposed approach in mitigating the adverse effects of sparsity on model performance on neuroimaging data. Notably, Global Random Mask outperformed DisPFL on lower sparsities, suggesting that in general global random masks might be more suitable for federated applications compared to even targeted unique local masks which DisPFL employs.

Dense baselines, such as FedAvg-FT and FedAvg, even while being *not sparse* and using full communication achieved comparable performances to NeuroSFL in the non-extreme sparsity region.

NeuroSFL even surpassed the performance of dense baselines at a sparsity level of 50%, highlighting the effectiveness of our proposed sparse method in optimizing model performance while reducing communication costs. Furthermore, Figure 2 (b) illustrates that the single global model trained with *NeuroSFL* demonstrated excellent performance for data within each site, emphasizing the model’s effectiveness in capturing site-specific characteristics while maintaining high accuracy.

Additionally, in Figure 3 (a), it is observed that the performance of local models trained with *NeuroSFL* remains consistently robust across non-IID states of local data, indicating the model’s versatility and reliability in various data distribution scenarios.

**Figure 3.**
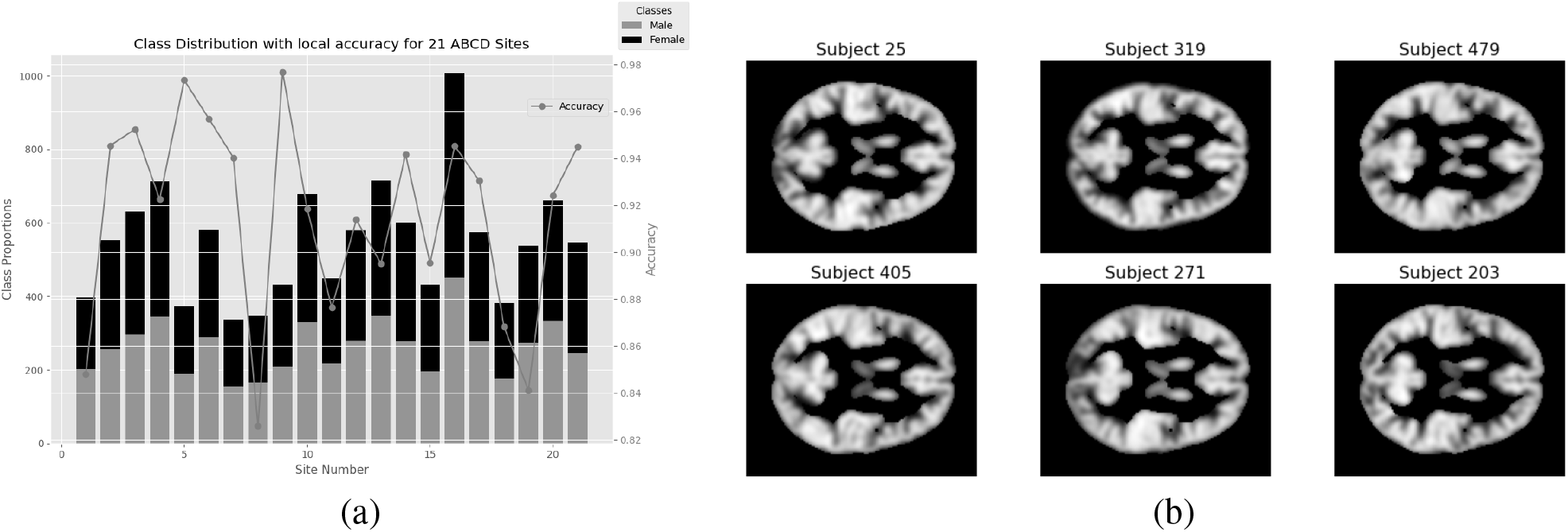
(a) Gender differences in each of the 21 ABCD sites along with the performance of the local models on each site with 50% sparsity. (b) Sample structural MRI brain image of brains used in gender classification.

#### 4.1.2 Effect of varying sparsity levels on non-IID data

##### Exploring the Impact of Varied Sparsity Levels on Non-IID Data for 10 Clients

In this subsection, we delve into the influence of changing sparsity levels on our model’s performance on non-IID data for 10 clients, employing the Dirichlet Distribution with alpha = 0.3 across various sparsity levels. Fig. 4 provides a visual representation of the accuracy versus sparsity relationship, showcasing the consistent accuracy achieved across different sparsity levels (Figure 4 (a)). Additionally, Figure 4 (b) illustrates the class distribution with Dir(0.3) for the ABCD dataset for 10 clients and their final local test accuracy. Following the Dirichlet partition, we have uneven data distribution, e.g. with site-6 having a total 5459 samples and sites-3 and 9 having only 8 and 5 training samples respectively.

**Figure 4.**
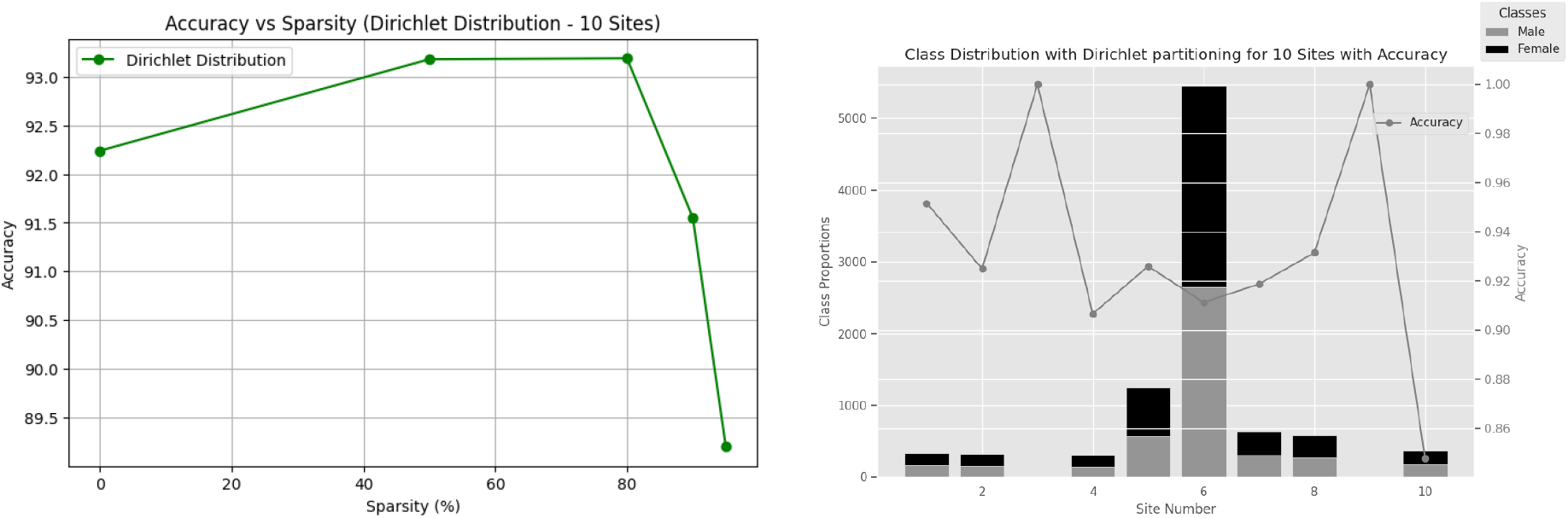
(a) Comparison of methods for gender classification using non-IID Dirichlet distribution with alpha=0.3 and varying sparsity levels. (b) Gender differences in each of the 10 ABCD sites along with the performance of the model.

Notably, our model demonstrated robust accuracy across different sparsity levels, ranging from 89.20% to 93.19%. These results underscore the resilience of our approach in maintaining high performance even under significant sparsity constraints. Importantly, when examining the local test accuracy across different clients, we observed consistently good performance regardless of the data differences. This resilience suggests that our model’s effectiveness extends beyond homogeneous datasets, making it suitable for deployment in federated learning scenarios with diverse client characteristics, specifically non-IID distribution.

##### Exploring the Impact of Varied Sparsity Levels on Non-IID Data for 30 Clients

We extend our analysis to incorporate 30 clients and explore the influence of changing sparsity levels on our model’s performance on non-IID data. We continue to employ the Dirichlet Distribution with alpha = 0.3 across various sparsity levels. Fig. 5 provides a visual representation of the accuracy versus sparsity relationship, showcasing the consistent accuracy achieved across different sparsity levels (fig. 5 (a)). Additionally, fig. 5 (b) illustrates the class distribution with Dir(0.3) for the ABCD dataset for 30 clients and their final local test accuracy. The results reveal a similar trend to the 10-client scenario, with the model achieving notable accuracy across varying sparsity levels. However, for the highly sparse condition of 95%, there is a slight drop in accuracy to 63.75%. This decline can be attributed to the increased difficulty for the model to generalize with such sparse data under the additional restriction of having larger clients. Despite this challenge, our model maintains its effectiveness across a diverse range of sparsity levels, indicating its potential for practical applications in federated learning scenarios with larger amount of client sites.

**Figure 5.**
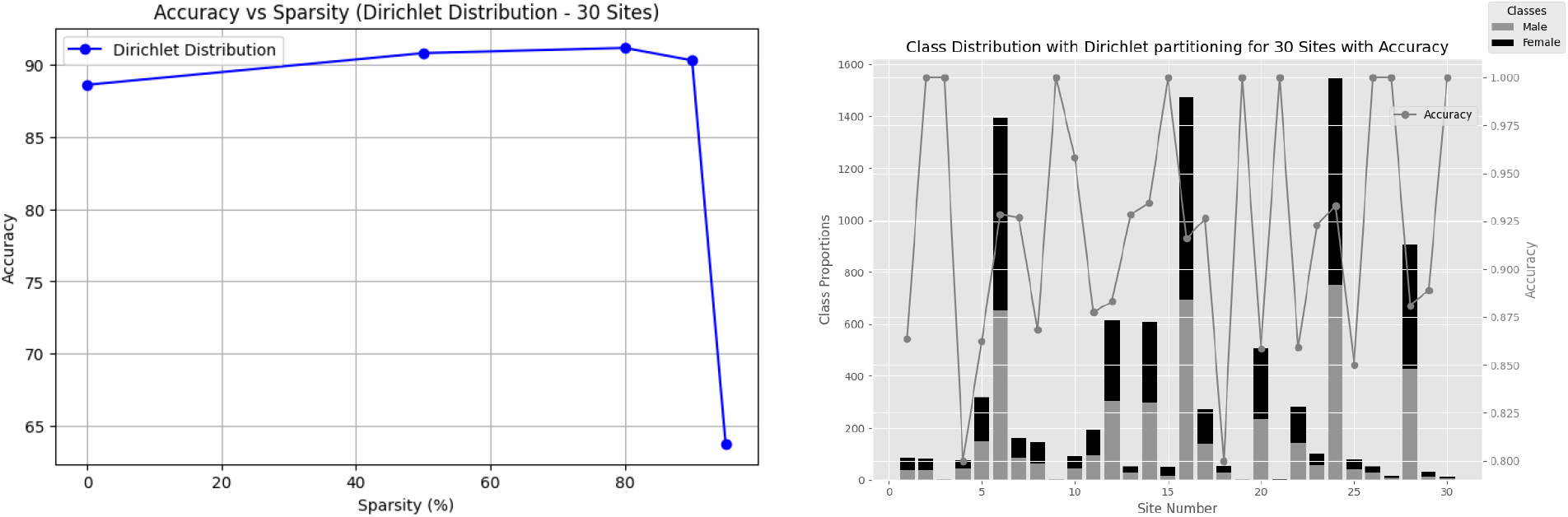
(a) Comparison of methods for gender classification using non-IID dirichlet distribution with alpha=0.3 and varying sparsity levels. (b) Gender differences in each of the 30 ABCD sites along with the performance of the model for 50% sps.

### 4.2 IterativeSNIP Performance

We evaluate the performance of IterSNIP and WeightedSNIP to explore their efficacy in sparse FL scenarios. Table 2 summarizes the accuracy results obtained at 50% sparsity for different iterations of IterSNIP and WeightedSNIP. IterSNIP, with varying numbers of iterations (1, 10, and 20), demonstrated consistent performance with increasing iterations, achieving accuracies of 92.4%, 91.82%, and 92.67%, respectively. These results suggest that utilizing multiple iterations to obtain SNIP masks does not necessarily enhance model performance in scenarios with sparsity constraints, and especially when used on neuroimaging data. This is a departure from the single-node case on natural image datasets such as CIFAR10 or CIFAR100 De Jorge et al. (2020). Similarly, WeightedSNIP, which incorporates weighted averages of saliency scores, achieved an accuracy of 92.10% and does not outperform the vanilla averaging technique. This proves that our model is robust enough to find a sparse mask, with minimal effect from the amount of data at each sites.

**Table 2.** Performance comparison of IterSNIP with different iterations and WeightedSNIP in terms of accuracy at sparsity of 50%.

| Method | Iterations | Accuracy (50% Sparsity) |
| --- | --- | --- |
| **IterSNIP** | 1 Iteration | 92.40% |
|  | 10 Iteration | 91.82% |
|  | 20 Iteration | 92.67% |
| **WeightedSNIP** | 1 Iteration | 92.10% |

### 4.3 Wall-time efficiency gains in the real world COINSTAC system

In our pursuit of optimizing federated learning methodologies, we make use of Coinstac Plis et al. (2016b), a cutting-edge open-source federated learning solution designed for collaborative neuroimaging endeavors at scale. Deployed in real-world scenarios, Coinstac embodies a paradigm shift in collaborative research, transcending traditional boundaries and fostering synergistic interactions among researchers worldwide. Coinstac’s architecture facilitates decentralized computations across a distributed network of geographically dispersed client nodes, seamlessly integrating diverse computational tasks while safeguarding data privacy through state-of-the-art differential privacy mechanisms. Having said this, our experiment leverages Coinstac’s robust infrastructure to benchmark our method against the standard dense FedAvg algorithm McMahan et al. (2017) within a practical real-world context. Our evaluation spans five diverse client locations, spanning from North Virginia to Frankfurt, each representing a distinct geographical node within Coinstac’s decentralized network. By meticulously assessing the mean communication time—reflecting the duration for the server model to aggregate all client weights during each communication round, we demonstrate the efficiency of our algorithm in optimizing federated learning workflows.

Our investigation encompasses five local client models, each featuring varying sizes or depths of ResNet architectures while maintaining a sparsity level of 90% across experiments. The ensuing comparative analysis, delineated in table 3, demonstrates the tangible speed enhancements achieved by our proposed methodology as compared to the standard FedAvg. Importantly, our results indicate that our approach consistently outperforms FedAvg across all ResNet architectures. For instance, in the case of ResNet32, our method achieves a communication time of 0.238 *±* 0.02 seconds, compared to 0.285 *±* 0.04 seconds for FedAvg, resulting in a speedup of 1.20x. This trend continues across deeper architectures, with our technique demonstrating significant improvements in communication efficiency. For instance, for ResNet110, our method achieves a remarkable speedup of 2.32x over FedAvg, showcasing its ability to handle complex models with greater efficiency. These empirical findings underscore the importance of sparse federated techniques like NeuroSFL, thereby propelling collaborative neuroimaging research to unprecedented heights.

**Table 3.** Comparison of communication time between FedAvg and NeuroSFL on Cifar10 for ResNet architectures of different depth.

| Architecture | Accuracy | Communication Time (s) |  | Speed up |
| --- | --- | --- | --- | --- |
|  |  | FedAvg | NeuroSFL |  |
| ResNet32 | 90.52% | $0.285 \pm 0.04$ | $0.238 \pm 0.02$ | 1.20x |
| ResNet44 | 89.65% | $0.409 \pm 0.06$ | $0.328 \pm 0.04$ | 1.24x |
| ResNet56 | 93.74% | $0.531 \pm 0.07$ | $0.407 \pm 0.06$ | 1.30x |
| ResNet110 | 93.25% | $1.812 \pm 0.33$ | $0.781 \pm 0.13$ | 2.32x |

### 4.4 Sparsity vs Accuracy Performance comparison

In this section we analyze and interpret the results from section 4. First, we probe the reasons behind the performance gains in comparison to a state of the art federated sparse learning method Dai et al. (2022).

In a specific comparison with DistPFL, we can see that NeuroSFL consistently performs better than DisPFL in a range of sparsities in the selected tasks. This is probably due to a better choice of the initial sparse sub-network using the importance criterion. Another difference is that, in DisPFL different local clients have different levels of sparsity and a final model averaging is done, where the final model becomes denser due to the union of many sparse subnetworks. We however retain the same mask in all the clients and start from the same initialization in all the clients, result in equivalent sparsity in all the clients; this also leaves open the potential of keeping sparse global models in a centralized FL setting.

## 5 CONCLUSION

In this paper, we elaborate a novel sparse decentralized federated learning approach that trains a sparse model efficiently for collaborative training on neuro-imaging data. By extending a gradient-based parameter importance criterion to the FL setting, we achieve reduced communication costs and better bandwidth in decentralized training. Our method leverages the nature of local data distribution, resulting in a client data-aware global sparse mask. This leads to savings in communication time and bandwidth during sparse training. We test our approach on the ABCD dataset and report improved performance compared to contemporary methods. Overall, our sparse FL technique enhances communication time, making it suitable for bandwidth-limited settings without compromising accuracy.

## AUTHOR CONTRIBUTIONS

BT and RO contributed equally in this work and share first authorship. EG assisted in manuscript preparation and helped in ideation. JL, VC and SP acted as advisors for the project and assisted in manuscript preparation and funded the project.

## REFERENCES

Abrol, A., Fu, Z., Salman, M., Silva, R., Du, Y., Plis, S., et al. (2021). Deep learning encodes robust discriminative neuroimaging representations to outperform standard machine learning. Nature Communications 12. doi:10.1038/s41467-020-20655-6

Arnob, S. Y., Ohib, R., Plis, S., and Precup, D. (2021). Single-shot pruning for offline reinforcement learning. arXiv preprint arXiv:2112.15579

Bibikar, S., Vikalo, H., Wang, Z., and Chen, X. (2022). Federated dynamic sparse training: Computing less, communicating less, yet learning better. In Proceedings of the AAAI Conference on Artificial Intelligence. vol. 36, 6080–6088

Bonawitz, K., Eichner, H., Grieskamp, W., Huba, D., Ingerman, A., Ivanov, V., et al. (2019). Towards federated learning at scale: System design. Proceedings of machine learning and systems 1, 374–388 [Dataset] Cerebras (2019). Wafer Scale Engine: Why We Need Big Chips for Deep Learning. https://cerebras.net/blog/cerebras-wafer-scale-engine-why-we-need-big-chips-for-deep-learning/

Chen, T., Frankle, J., Chang, S., Liu, S., Zhang, Y., Wang, Z., et al. (2020). The lottery ticket hypothesis for pre-trained bert networks. Advances in neural information processing systems 33, 15834–15846

Cheng, G., Chadha, K., and Duchi, J. (2021). Fine-tuning is fine in federated learning. arXiv preprint arXiv:2108.07313

Dai, R., Shen, L., He, F., Tian, X., and Tao, D. (2022). Dispfl: Towards communication-efficient personalized federated learning via decentralized sparse training. arXiv preprint arXiv:2206.00187

De Jorge, P., Sanyal, A., Behl, H. S., Torr, P. H., Rogez, G., and Dokania, P. K. (2020). Progressive skeletonization: Trimming more fat from a network at initialization. arXiv preprint arXiv:2006.09081

Elsen, E., Dukhan, M., Gale, T., and Simonyan, K. (2020). Fast sparse convnets. In Proceedings of the IEEE/CVF conference on computer vision and pattern recognition. 14629–14638

Evci, U., Gale, T., Menick, J., Castro, P. S., and Elsen, E. (2020). Rigging the lottery: Making all tickets winners. In International Conference on Machine Learning (PMLR), 2943–2952

Frankle, J. and Carbin, M. (2019). The lottery ticket hypothesis: Finding sparse, trainable neural networks. In 7th International Conference on Learning Representations, ICLR 2019, New Orleans, LA, USA, May 6-9, 2019

Frankle, J., Dziugaite, G. K., Roy, D., and Carbin, M. (2021). Pruning neural networks at initialization: Why are we missing the mark? In 9th International Conference on Learning Representations, ICLR 2021, Virtual Event, Austria, May 3-7, 2021

Garavan, H., Bartsch, H., Conway, K., Decastro, A., Goldstein, R., Heeringa, S., et al. (2018). Recruiting the abcd sample: Design considerations and procedures. Developmental cognitive neuroscience 32, 16–22

Han, S., Mao, H., and Dally, W. J. (2015). Deep compression: Compressing deep neural networks with pruning, trained quantization and huffman coding. arXiv preprint arXiv:1510.00149

Hsu, T.-M. H., Qi, H., and Brown, M. (2019). Measuring the effects of non-identical data distribution for federated visual classification. arXiv preprint arXiv:1909.06335

Huang, T., Liu, S., Shen, L., He, F., Lin, W., and Tao, D. (2022). Achieving personalized federated learning with sparse local models. arXiv preprint arXiv:2201.11380

Jiang, Y., Wang, S., Valls, V., Ko, B. J., Lee, W.-H., Leung, K. K., et al. (2022). Model pruning enables efficient federated learning on edge devices. IEEE Transactions on Neural Networks and Learning Systems

Konečný, J., McMahan, H. B., Ramage, D., and Richtárik, P. (2016). Federated optimization: Distributed machine learning for on-device intelligence. arXiv preprint arXiv:1610.02527

Krizhevsky, A., Sutskever, I., and Hinton, G. E. (2012). Imagenet classification with deep convolutional neural networks. In Advances in Neural Information Processing Systems, eds. F. Pereira, C. Burges, L. Bottou, and K. Weinberger (Curran Associates, Inc.), vol. 25

Kulkarni, V., Kulkarni, M., and Pant, A. (2020). Survey of personalization techniques for federated learning. In 2020 Fourth World Conference on Smart Trends in Systems, Security and Sustainability (WorldS4) (IEEE), 794–797

Laird, A. R. (2021). Large, open datasets for human connectomics research: Considerations for reproducible and responsible data use. NeuroImage 244, 118579

Landis, D., Courtney, W., Dieringer, C., Kelly, R., King, M., Miller, B., et al. (2016). Coins data exchange: An open platform for compiling, curating, and disseminating neuroimaging data. NeuroImage 124, 1084–1088

Lee, N., Ajanthan, T., and Torr, P. H. (2018). Snip: Single-shot network pruning based on connection sensitivity. arXiv preprint arXiv:1810.02340

Li, A., Sun, J., Wang, B., Duan, L., Li, S., Chen, Y., et al. (2020a). Lotteryfl: Personalized and communication-efficient federated learning with lottery ticket hypothesis on non-iid datasets. arXiv preprint arXiv:2008.03371

Li, A., Sun, J., Zeng, X., Zhang, M., Li, H., and Chen, Y. (2021). Fedmask: Joint computation and communication-efficient personalized federated learning via heterogeneous masking. In Proceedings of the 19th ACM Conference on Embedded Networked Sensor Systems. 42–55

Li, W., Milletarì, F., Xu, D., Rieke, N., Hancox, J., Zhu, W., et al. (2019). Privacy-preserving federated brain tumour segmentation. In Machine Learning in Medical Imaging: 10th International Workshop, MLMI 2019, Held in Conjunction with MICCAI 2019, Shenzhen, China, October 13, 2019, Proceedings 10 (Springer), 133–141

Li, X., Gu, Y., Dvornek, N., Staib, L. H., Ventola, P., and Duncan, J. S. (2020b). Multi-site fmri analysis using privacy-preserving federated learning and domain adaptation: Abide results. Medical Image Analysis 65, 101765

Ma, R., Miao, J., Niu, L., and Zhang, P. (2019). Transformed ℓ1 regularization for learning sparse deep neural networks. Neural Networks 119, 286–298

McMahan, B., Moore, E., Ramage, D., Hampson, S., and y Arcas, B. A. (2017). Communication-efficient learning of deep networks from decentralized data. In Artificial intelligence and statistics (PMLR), 1273–1282

[Dataset] McMahan, B. and Ramage, D. (2017). Federated learning: Collaborative machine learning without centralized training data

McMahan, H. B., Moore, E., Ramage, D., Hampson, S., and y Arcas, B. A. (2016). Communication-efficient learning of deep networks from decentralized data. In International Conference on Artificial Intelligence and Statistics

Milham, M. P., Craddock, R. C., Son, J. J., Fleischmann, M., Clucas, J., Xu, H., et al. (2018). Assessment of the impact of shared brain imaging data on the scientific literature. Nature Communications 9, 2818

Ming, J., Verner, E., Sarwate, A., Kelly, R., Reed, C., Kahleck, T., et al. (2017). Coinstac: Decentralizing the future of brain imaging analysis. F1000Research 6

Mozer, M. C. and Smolensky, P. (1988). Skeletonization: A technique for trimming the fat from a network via relevance assessment. Advances in neural information processing systems 1

Ohib, R., Gillis, N., Dalmasso, N., Shah, S., Potluru, V. K., and Plis, S. (2022). Explicit group sparse projection with applications to deep learning and NMF. Transactions on Machine Learning Research

Ohib, R., Thapaliya, B., Gaggenapalli, P., Liu, J., Calhoun, V., and Plis, S. (2023). Salientgrads: Sparse models for communication efficient and data aware distributed federated training. arXiv preprint arXiv:2304.07488

Plis, S. M., Sarwate, A. D., Wood, D., Dieringer, C., Landis, D., Reed, C., et al. (2016a). Coinstac: a privacy enabled model and prototype for leveraging and processing decentralized brain imaging data. Frontiers in neuroscience 10, 204805

Plis, S. M., Sarwate, A. D., Wood, D., Dieringer, C., Landis, D., Reed, C., et al. (2016b). COINSTAC: A privacy enabled model and prototype for leveraging and processing decentralized brain imaging data. Frontiers in Neuroscience 10. doi:10.3389/fnins.2016.00365

Poldrack, R. A., Barch, D. M., Mitchell, J. P., Wager, T. D., Wagner, A. D., Devlin, J. T., et al. (2013). Toward open sharing of task-based fmri data: the openfmri project. Frontiers in neuroinformatics 7, 12

[Dataset] Pool, J., Sawarkar, A., and Rodge, J. (2021). Accelerating Inference with Sparsity Using the NVIDIA Ampere Architecture and NVIDIA TensorRT. https://developer.nvidia.com/blog/accelerating-inference-with-sparsity-using-ampere-and-tensorrt/

Qiu, X., Fernandez-Marques, J., Gusmao, P. P., Gao, Y., Parcollet, T., and Lane, N. D. (2022). Zerofl: Efficient on-device training for federated learning with local sparsity. arXiv preprint arXiv:2208.02507

Renda, A., Frankle, J., and Carbin, M. (2020). Comparing rewinding and fine-tuning in neural network pruning. arXiv preprint arXiv:2003.02389

Rootes-Murdy, K., Gazula, H., Verner, E., Kelly, R., DeRamus, T., Plis, S., et al. (2022). Federated analysis of neuroimaging data: a review of the field. Neuroinformatics, 1–14

Roy, A. G., Siddiqui, S., Pölsterl, S., Navab, N., and Wachinger, C. (2019). Braintorrent: A peer-to-peer environment for decentralized federated learning. arXiv preprint arXiv:1905.06731

Sheller, M. J., Reina, G. A., Edwards, B., Martin, J., and Bakas, S. (2019). Multi-institutional deep learning modeling without sharing patient data: A feasibility study on brain tumor segmentation. In Brainlesion: Glioma, Multiple Sclerosis, Stroke and Traumatic Brain Injuries: 4th International Workshop, BrainLes 2018, Held in Conjunction with MICCAI 2018, Granada, Spain, September 16, 2018, Revised Selected Papers, Part I 4 (Springer), 92–104

Silva, S., Gutman, B. A., Romero, E., Thompson, P. M., Altmann, A., and Lorenzi, M. (2019). Federated learning in distributed medical databases: Meta-analysis of large-scale subcortical brain data. In 2019 IEEE 16th international symposium on biomedical imaging (ISBI 2019) (IEEE), 270–274

Sokar, G., Mocanu, E., Mocanu, D. C., Pechenizkiy, M., and Stone, P. (2021). Dynamic sparse training for deep reinforcement learning. arXiv preprint arXiv:2106.04217

Tanaka, H., Kunin, D., Yamins, D. L., and Ganguli, S. (2020). Pruning neural networks without any data by iteratively conserving synaptic flow. In Advances in Neural Information Processing Systems 33: Annual Conference on Neural Information Processing Systems 2020, NeurIPS 2020, December 6-12, 2020, virtual, eds. H. Larochelle, M. Ranzato, R. Hadsell, M. Balcan, and H. Lin

Tenopir, C., Allard, S., Douglass, K., Aydinoglu, A. U., Wu, L., Read, E., et al. (2011). Data sharing by scientists: practices and perceptions. PloS one 6, e21101

Wang, C., Zhang, G., and Grosse, R. B. (2020). Picking winning tickets before training by preserving gradient flow. In 8th International Conference on Learning Representations, ICLR 2020, Addis Ababa, Ethiopia, April 26-30, 2020

Yang, H., Wen, W., and Li, H. (2019). Deephoyer: Learning sparser neural network with differentiable scale-invariant sparsity measures. arXiv preprint arXiv:1908.09979

Zhao, Y., Li, M., Lai, L., Suda, N., Civin, D., and Chandra, V. (2018). Federated learning with non-iid data. arXiv preprint arXiv:1806.00582

Zhu, M. and Gupta, S. (2018). To prune, or not to prune: Exploring the efficacy of pruning for model compression. In 6th International Conference on Learning Representations, ICLR 2018, Vancouver, BC, Canada, April 30 - May 3, 2018, Workshop Track Proceedings

